# A developmental timer coordinates organism-wide microRNA transcription

**DOI:** 10.64898/2026.01.21.700890

**Authors:** Peipei Wu, Jing Wang, Brett Pryor, Isabella Valentino, David F. Ritter, Kaiser Loel, Justin Kinney, Sevinc Ercan, Leemor Joshua-Tor, Christopher M. Hammell

## Abstract

The development of distinct tissues must be precisely coordinated to ensure that growth and cell fate transitions occur in the correct temporal order across the organism, yet the mechanisms that coordinate these timing events remain unclear. In *Caenorhabditis elegans*, stage-specific cell fate transitions are driven by pulsatile transcription of heterochronic microRNAs, but the source of these rhythms has been unknown. Here, we identify a developmental timer composed of the transcription factor MYRF-1 and the PERIOD-like repressor LIN-42 that operates in all somatic cells. MYRF-1 binds conserved regulatory elements upstream of heterochronic microRNA genes and drives synchronized, once-per-stage transcriptional pulses across tissues, while concurrently activating *lin-42* expression. Newly synthesized LIN-42 directly associates with MYRF-1, limiting its nuclear residence and transcriptional activity to constrain the amplitude and duration of each transcriptional burst. This reciprocal transcriptional/translational feedback loop generates organism-wide, phase-locked microRNA expression, coupling tissue-specific development to organismal growth through a shared timing mechanism.

## INTRODUCTION

During animal development, various cell types must acquire specialized identities while remaining coordinated with organismal growth. Such coordination depends not only on correct fate specification but also on the alignment of fate transitions across tissues, ensuring that proliferation, differentiation, and morphogenesis unfold in the proper relative sequence. Yet the mechanisms that generate this organism-wide temporal coherence remain unclear. It is unknown whether distinct cell lineages share a unified temporal program for measuring developmental time, or whether timing instead emerges from lineage-intrinsic regulatory architectures that independently encode the timing of fate transitions.

In *C. elegans*, the sequence of temporal patterning is controlled by a conserved heterochronic gene network, centered on key microRNAs (miRNAs) that enforce synchronous, stage-specific cell-fate transitions across tissues (*1-7*). These miRNAs repress transcription factors and RNA-binding proteins that normally coordinate stage-specific proliferation, differentiation, and morphogenesis, and block the precocious onset of later developmental programs (*8, 9*). Mutations in heterochronic miRNAs disrupt the progression of temporal events throughout the organism, highlighting their global role in developmental transitions(*1, 2, 5*). Interestingly, heterochronic miRNAs are transcribed in sharp, once-per-larval-stage bursts in proliferating blast cells, differentiating epithelia, intestinal cells, glia, and post-mitotic neurons(*10-13*). Although their rhythmic expression parallels the oscillatory transcription of many protein-coding genes, the two processes operate on distinct organizational scales: oscillatory mRNAs display a wide range of tissue-specific expression windows within each larval stage(*14, 15*), whereas heterochronic miRNAs are transcribed in a shared phase of each larval stage across most somatic tissues(*12, 16*).

This synchrony raises a central question: are heterochronic miRNAs regulated across somatic cells by lineage-specific transcriptional programs that require unique transcription factor networks to repeatedly converge on a common timing, or does a shared molecular mechanism enforce organism-wide coherence? Several cell-type-specific transcription factors modulate miRNA dynamics in individual tissues(*13, 17, 18*), yet none have been shown to drive the global, once-per-stage transcriptional bursts necessary for system-wide coordination. In contrast, both genetic analyses and direct measurements of miRNA transcriptional dynamics in animals harboring mutations in *lin-42*, encoding the nematode ortholog of the circadian regulator *Period*, suggest that LIN-42 acts as a widespread transcriptional repressor of miRNA genes whose own expression is pulsatile and phase-coherent across somatic tissues(*12, 13, 19-22*). These properties indicate that LIN-42 acts as a key regulator of organism-wide timing and heterochronic miRNA transcription, even though *C. elegans* does not encode orthologs of the transcription factors that are usually repressed by Period in other systems (such as Clock and Bmal in mice and humans, or Clock and Cycle in *Drosophila*). Together, these observations highlight a critical gap in understanding: although LIN-42 clearly imposes soma-wide temporal coordination of heterochronic miRNA transcription, the underlying mechanism—and the transcription factors whose activities LIN-42 antagonizes to generate these organism-wide transcriptional pulses—remain unknown.

### MYRF-1 regulates the expression of heterochronic miRNAs and *lin-42*

The *myrf-1* gene encodes an essential homotrimeric transcription factor that accumulates rhythmically, once per larval stage, in all somatic tissues (fig. S1)(*10, 14, 15, 23, 24*). Similar to its mammalian ortholog, MYRF-1 is synthesized as a full-length membrane-associated precursor that undergoes self-cleavage to release an N-terminal nuclear domain (MYRF-1(ND))(*23*). Genetic analysis places MYRF-1 in early larval development and upstream of the transcriptional activation of the heterochronic miRNA *lin-4*: loss of *myrf-1* function strongly reduces activity of a *lin-4* transcriptional reporter and decreases mature *lin-4* RNA levels(*24*). Notably, the onset of MYRF-1 nuclear accumulation coincides with the initiation of *lin-4* transcription, suggesting direct transcriptional control(*24*).

To define the genomic targets of MYRF-1, we performed ChIP–seq on staged animals expressing an endogenously tagged GFP::MYRF-1 fusion at peak nuclear abundance during the L1 stage. This analysis identified ∼1,000 high-confidence MYRF-1 binding sites, predominantly located within 3 kb of transcription start sites of potential MYRF-1 target genes (Table S1; fig. S1e). Target genes were significantly enriched for regulators of temporal patterning, larval development, and ribosome biogenesis (fig. S1f). Strikingly, MYRF-1 binding was strongly enriched at conserved regulatory regions upstream of all heterochronic miRNA genes (Fig. 1a), as well as at promoters of key oscillatory regulators, including *lin-42*, (Fig. 1b), *myrf-1* itself, and multiple genes required for molting (*nhr-23, grh-1*, and *mab-10*) (fig. S1x). Motif analysis of sequences over-represented in MYRF-1 peaks revealed a conserved GA-rich sequence composed of three elements with defined orientation and spacing (Fig. 1c), consistent with multimeric DNA binding. In agreement with this model, a recombinant MYRF-1 protein fragment (residues 1-483) that corresponds to the MYRF-1(ND) binds specifically to a 44-bp element within the proximal *lin-4* regulatory region (proxB) that conforms to the MYRF-1 consensus motif (Figure 1a, c, d, and e). This MYRF-1:proxB binary complex forms stably *in vitro* and elutes at a volume consistent with a trimer on DNA according to size exclusion chromatography and mass photometry (Fig. 1 f and fig. S2).

**Figure 1.**
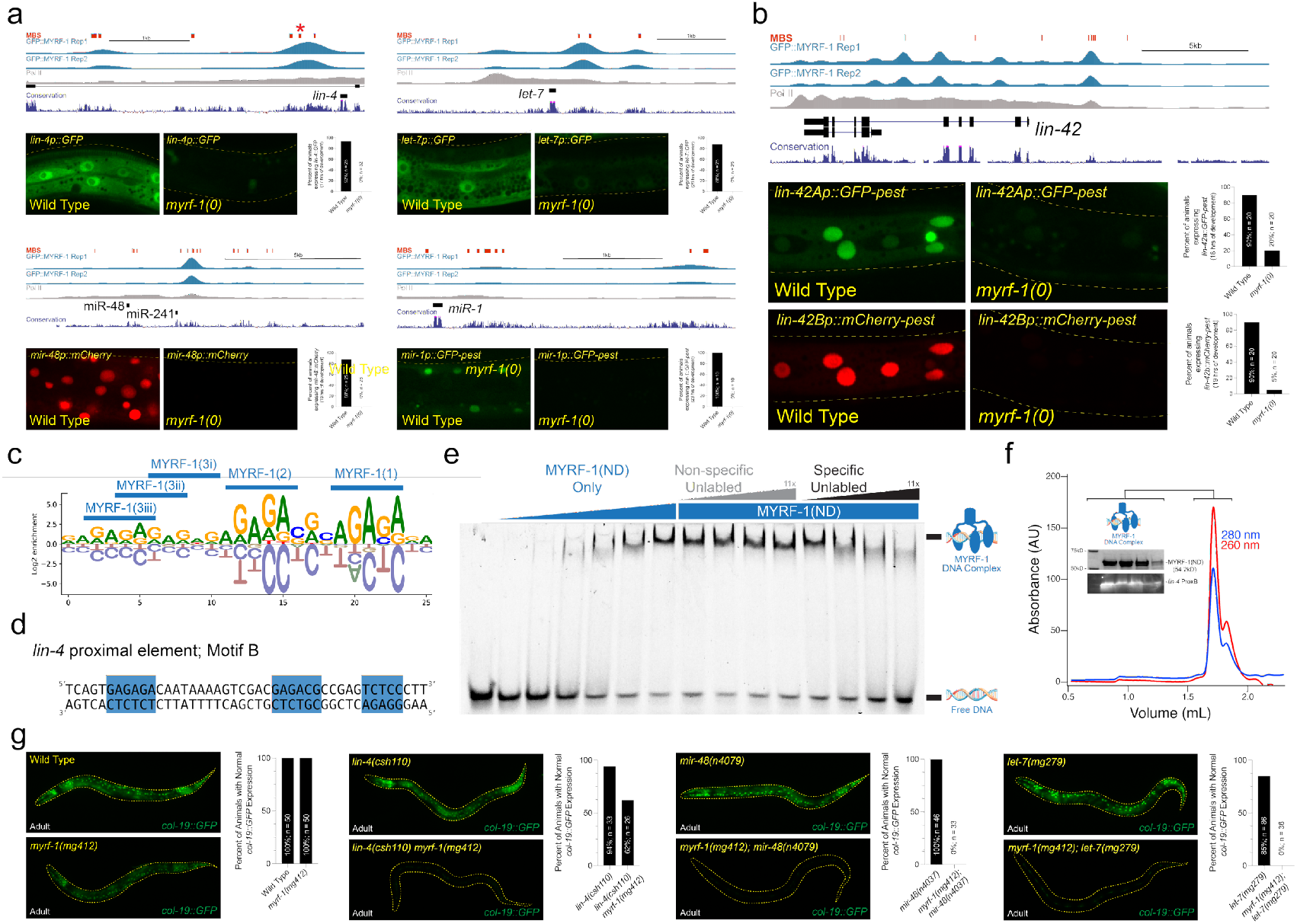
MYRF-1 controls temporal patterning by binding to sequences upstream of heterochronic miRNAs and the *lin-42* gene. **a and b**, MYRF-1 binding sites are found in the putative regulatory regions of cyclically expressed miRNAs and the *lin-42*. Transcriptional reporters of these MYRF-1 target genes require *myrf-1* for expression. **c and d**, Motif analysis of sequences found within MYRF-1 binding sites identifies a consensus motif harboring a repetitive GA-rich sequence. Sequences that conform to this consensus are indicated in panels a and b. An example of one of these binding sites from the *lin-4* regulatory region, *lin-4* proximal element motif B, is shown in panel d. The location of this putative MYRF-1 binding site is indicated with an asterisk in panel a. **e**, Recombinant MYRF-1(ND) binds specifically to the *lin-4* proximal element motif B DNA fragment. **f**, Recombinant MYRF-1(ND) forms a trimeric complex that co-purifies with a *lin-4* proximal element motif B DNA (ProxB) during size exclusion chromatography. **g**, A hypomorphic allele of *myrf-1, myrf-1(mg412*), does not exhibit defects in the expression of an adult-specific transcriptional reporter (*col-19p::GFP*), but strongly enhances the temporal patterning defects associated with mutations in heterochronic miRNAs.

To define the requirement for MYRF-1 in developmental gene activation, we analyzed transcriptional reporters in *myrf-1* null mutants that arrest during the L1 molt (*25*). In wild-type L1 larvae, heterochronic miRNA transcriptional reporters—including *lin-4* and *let-7* family members—were robustly expressed, whereas all were completely silent in *myrf-1(0*) animals (Fig. 1a). In contrast, among oscillatory protein-coding targets examined, only *lin-42* expression depended on MYRF-1, while other molting regulators remained active (Fig. 1b; fig. S1d). These results establish MYRF-1 as a direct transcriptional activator of heterochronic miRNAs and *lin-42* and reveal that rhythmic gene expression during larval development is generated by at least two mechanistically distinct regulatory systems.

We reasoned that if MYRF-1 controls heterochronic miRNA expression at larval stages after the L1, then partial loss of MYRF-1 activity should enhance phenotypes associated with miRNA mutants that function in the L2-adult stages of development. This type of genetic interaction would lead to the reiteration of distinct cell-fate specification events at later larval stages. The *myrf-1(mg412*) allele alters amino acids near the predicted DNA-binding domain of MYRF-1, leading to an inappropriate reiteration of larval molting cycles in adults(*25*). Importantly, *myrf-1(mg412*) mutant animals do not exhibit heterochronic phenotypes and properly express an adult-specific *col-19p::GFP* reporter (Fig.1g; Table 1). However, combining *myrf-1(mg412*) with a hypomorphic *lin-4* allele (*lin-4(csh110)*) produced highly penetrant synthetic retarded phenotypes, including reduced *col-19p::GFP* expression in adult-stage animals and loss of alae structures on adult cuticles (Fig. 1g; Table 1). *myrf-1(mg412*) also strongly enhanced defects in a *mir-48* deletion mutant, *mir-48(n4097*), resulting in the failure of adult epidermal differentiation, reiteration of L2-stage cell division programs during L3 stage, and defective production of adult-specific alae structures (Fig. 1g; Table 1 fig. Sx).

**Table 1.** C. *elegans myrf* genes genetically interacts with multiple heterochronic mutants to control temporal patterning.

| Genotype<sup>b</sup> | Percent of animals with Adult-specific alae formation<sup>a</sup> |  |  |  |  |  |  |  |
| --- | --- | --- | --- | --- | --- | --- | --- | --- |
|  | L4 stage |  |  |  | Adult stage |  |  |  |
|  | none | gapped | full | n = | none | gapped | full | n = |
| wild type @20°C | 100 | 0 | 0 | 20 | 0 | 0 | 100 | 20 |
| wild type @15°C | 100 | 0 | 0 | 20 | 0 | 0 | 100 | 20 |
| *myrf-1(mg412)* @20°C | 100 | 0 | 0 | 20 | 0 | 0 | 100 | 22 |
| *myrf-1(mg412)* @15°C | 100 | 0 | 0 | 20 | 0 | 0 | 100 | 22 |
| *myrf-2(gk669)* | 100 | 0 | 0 | 20 | 0 | 0 | 100 | 20 |
| *myrf-1(mg412); myrf-2(gk669)* | 100 | 0 | 0 | 20 | 0 | 0 | 100 | 20 |
| *lin-42(n1089)* | 9 | 30 | 61 | 23 | 0 | 35 | 65 | 20 |
| *lin-42(n1089) myrf-1(mg412)* | 100 | 0 | 0 | 20 | 0 | 5 | 95 | 21 |
| *lin-42(n1089); myrf-2(gk669)* | 30 | 65 | 5 | 20 | 0 | 59 | 41 | 22 |
| *lin-42(n1089) myrf-1(mg412); myrf-2(gk669)* | 100 | 0 | 0 | 22 | 0 | 0 | 100 | 20 |
| *lin-42(ok2385)* | 29 | 38 | 33 | 24 | 0 | 33 | 67 | 21 |
| *lin-42(ok2385) myrf-1(mg412)* | 90 | 0 | 10 | 20 | 0 | 0 | 100 | 25 |
| *lin-46(ma164)* @20°C | - | - | - | - | 0 | 0 | 100 | 21 |
| *lin-46(ma164)* @15°C | - | - | - | - | 0 | 70 | 30 | 20 |
| *myrf-1(mg412); lin-46(ma164)* @20°C | - | - | - | - | 0 | 0 | 100 | 20 |
| *myrf-1(mg412); lin-46(ma164)* @15°C | - | - | - | - | 9 | 91 | 0 | 22 |
| *lin-4(csh104)* | - | - | - | - | 100 | 0 | 0 | 21 |
| *lin-4(csh110)* | - | - | - | - | 4 | 50 | 46 | 24 |
| *lin-4(csh111)* | - | - | - | - | 0 | 29 | 71 | 21 |
| *lin-4(csh110 csh111)* | - | - | - | - | 90 | 10 | 0 | 21 |
| *myrf-1(mg412) lin-4(csh110)* | - | - | - | - | 61 | 26 | 13 | 23 |
| *mir-48(n4097)* | - | - | - | - | 0 | 0 | 100 | 20 |
| *myrf-1(mg412); mir-48(n4097)* | - | - | - | - | 77 | 23 | 0 | 22 |
| *let-7(mg285)* | - | - | - | - | 0 | 0 | 100 | 20 |
| *myrf-1(mg412); let-7(mg285)* | - | - | - | - | 42 | 58 | 0 | 24 |
| *lin-42(csh86 (ΔMBD1))* | 100 | 0 | 0 | 20 | 0 | 0 | 100 | 24 |
| *lin-42(csh83 (ΔMBD2))* | 100 | 0 | 0 | 21 | 0 | 0 | 100 | 20 |
| *lin-42(csh83 csh86 (ΔMBD1+2))* | 22 | 46 | 32 | 22 | 0 | 55 | 45 | 20 |

<sup>a</sup>Presence and quality (Gapped or Complete) of cuticular alae structures were assayed by Normarski DIC optics. Only one side of each animal was scored.
<sup>b</sup>Animals contain *mals105*, which expresses an adult-specific, *col-19p::GFP* reporter integrated on chromosome V.

Consistent with a broad role in temporal progression, even in late temporal cell fate transitions, *myrf-1(mg412*) enhanced defects in a *let-7* hypomorphic mutant (*let-7(mg279)*) (Figure 1g; Table 1) and exhibited stage-specific synthetic phenotypes when combined with mutations in factors that prime heterochronic miRNA transcription (*blmp-1*) or modulate post-translational repression of HBL-1 (*lin-46*) (fig. S1h–j). Importantly, RNAi-mediated depletion of *hbl-1* fully suppressed these late-stage synthetic phenotypes, indicating that MYRF-1 promotes temporal progression primarily by enabling heterochronic miRNA–mediated repression of temporal identity genes (fig. S1x). Together, these results demonstrate that MYRF-1 acts broadly and cooperatively within the heterochronic pathway to ensure correct temporal progression, principally by driving transcription of heterochronic miRNAs.

### Chromatin accessibility dictates MYRF-1 functionality in diverse cell lineages

To determine whether MYRF-1 binding upstream of *lin-4* is necessary for correct temporal patterning amongst diverse cell lineages, we integrated our ChIP-seq-defined MYRF-1 binding site data with cell-type–resolved chromatin accessibility maps derived from single-cell ATAC-seq experiments(*26*). Both MYRF-1 binding regions upstream of *lin-4* are within accessible chromatin in hypodermal lineages at the L2 stage (Fig. 2a). In contrast, only the proximal MYRF-1 binding region is accessible in neuronal lineages (Fig. 2a), indicating that MYRF-1-dependent regulation of *lin-4* may be limited by lineage-specific chromatin structure. The functional contribution of these elements was then probed genetically by using CRISPR/Cas-9 genome editing to delete either the distal or proximal MYRF-1 binding regions individually or in combination in the endogenous context (Fig. 2a). Deletion of either element alone produced minimal developmental defects and regular expression of temporal patterning reporters in hypodermal tissues (Fig. 2b; Table 1). By contrast, simultaneous deletion of both regions resulted in highly penetrant heterochronic phenotypes that closely phenocopied *lin-4(0*) mutants (Fig. 2b; Table 1; Table S2), demonstrating that MYRF-1 binding to either site is sufficient to support normal hypodermal temporal patterning and that both accessible enhancers bound by MYRF-1 are functionally redundant in developing nematode skin cells.

**Figure 2.**
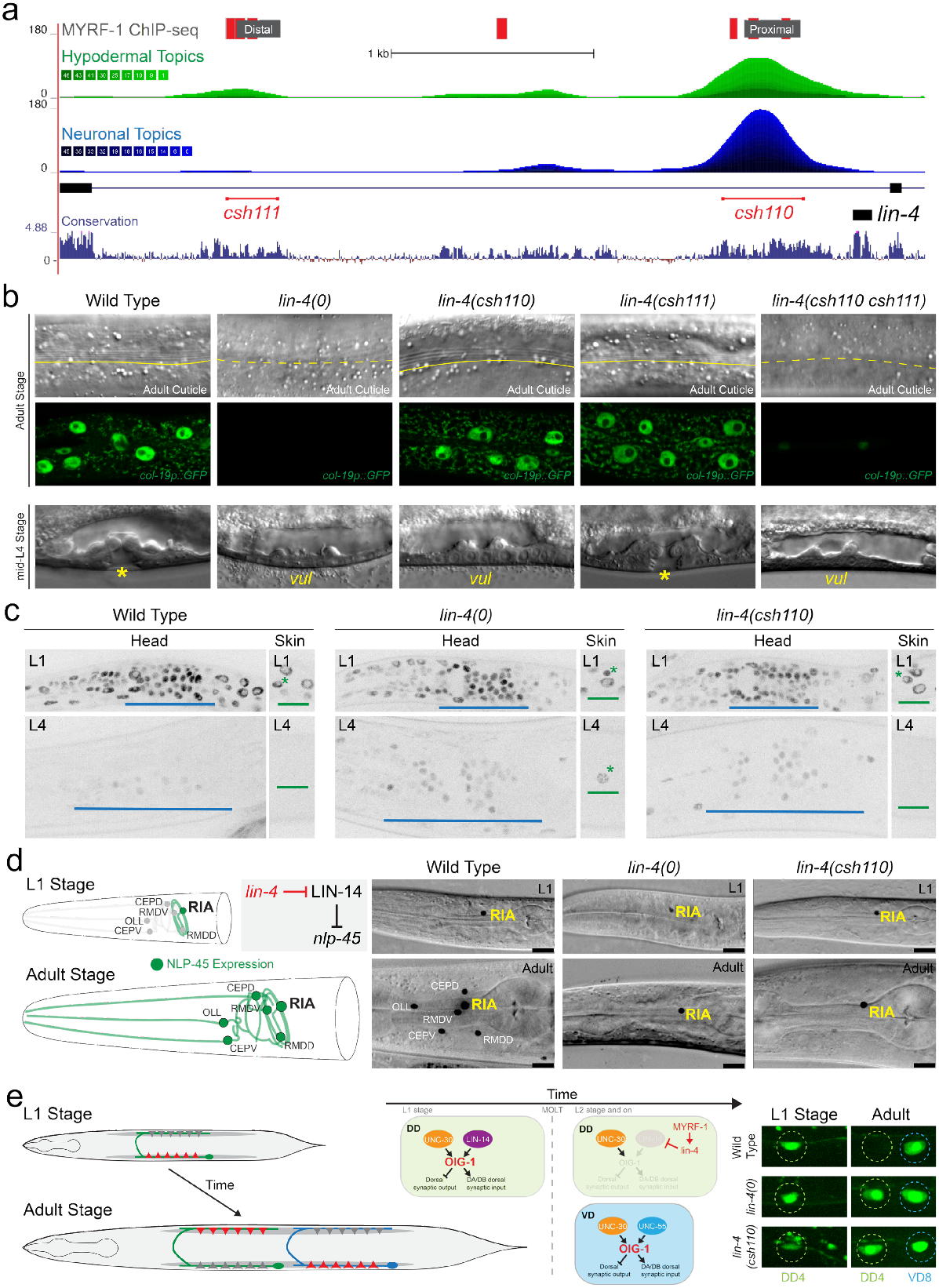
Differential requirements for MYRF-1 binding sites at the *lin-4* locus across tissues. **a**, Schematic of the *lin-4* locus showing GFP::MYRF-1 binding sites (grey) and predicted MYRF-1 consensus motifs (red). Genome browser tracks display chromatin accessibility in hypodermal (green) and neuronal (blue) cell types from L2-stage scATAC-seq data. Conservation of the *lin-4* locus across nematode species is shown below, along with the positions of the *lin-4(csh111*) and *lin-4(csh110*) alleles, which delete clusters of MYRF-1 binding sites. **b**, Representative micrographs showing adult cuticle and *col-19p::GFP* expression, and L4-stage vulval morphology, in animals of the indicated genotypes. **c**, Expression dynamics of LIN-14::GFP in wild-type and *lin-4* mutant animals. Blue bars denote LIN-14::GFP expression in head neuronal ganglia; green bars denote expression in hypodermal tissues. Green asterisks mark lateral seam cells expressing LIN-14::GFP at the indicated stages. **d**, Schematic and representative images of *nlp-45* expression in head ganglion neurons. Following LIN-14 downregulation at the end of L1, *nlp-45* expression expands to additional sensory and motor neurons. Images show *nlp-45::T2A::GFP::H2B* reporter expression in staged animals of the indicated genotypes. Phenotypes are fully penetrant in late L4 (0% of wild type (n = 25), 100% of *lin-4(0*) (n = 25), and 100% of *lin-4(csh110*) (n = 22). **e**, Persistent expression of an *oig-1::GFP* transcriptional reporter in DD neurons of *lin-4* mutants. In *lin-4(0*) and *lin-4(csh110*) mutants, an *oig-1p::GFP* transcriptional reporter expression persists into late L4 (0/25 wild type, 20/20 *lin-4(0*), 22/23 *lin-4(csh110)*).

Despite this redundancy in the skin, deletion of the proximal element (*lin-4(csh110)*) caused a fully penetrant egg-laying defective phenotype (n = 120) that was absent in wild-type (n = 60) or in animals lacking the distal enhancer element, *lin-4(csh111*) (n = 100), revealing a lineage-specific requirement for MYRF-1 input. This defect correlated with failure of vulval precursor cell (VPC) maturation (Fig. 2b; Table S3). Moreover, even in *csh110* animals with grossly normal VPC cell lineages, axons of the Hermaphrodite-Specific Neuron (HSN) that promote egg laying in adulthood failed to extend and innervate adult vulval structures, closely resembling the neuronal defects observed in *lin-4(0*) mutants (fig. SX)(*27*). These findings suggest that neuronal and vulval lineages are more sensitive to loss of the proximal MYRF-1 binding region than hypodermal tissues.

To directly examine the consequences of these cis-regulatory perturbations on *lin-4* function, we monitored the temporal expression dynamics of its direct target, the transcription factor LIN-14. In wild-type animals, *lin-4* expression during mid-L1 leads to repression of LIN-14 across somatic tissues (Fig. 2c)(*2, 3, 28*). As expected, *lin-4(0*) mutants fail to repress LIN-14, resulting in persistent LIN-14::GFP expression in both hypodermal and neuronal lineages (Fig. 2c)(*29*). In *lin-4(csh110*) animals, LIN-14::GFP was properly downregulated in hypodermal cells by the end of L1, but persisted at high levels in neurons, closely resembling the neuronal levels of LIN-14::GFP in *lin-4(0*) mutant neurons (Fig. 2c). By contrast, LIN-14 regulation in *lin-4(csh111*) mutants was indistinguishable from wild type. Consistent with these lineage-specific defects in LIN-14 regulation, *lin-4(csh110*) animals showed pronounced temporal mosaicism: hypodermal cells executed normal temporal programs and matured to nearly normal adult tissues, whereas multiple neuronal cell types displayed fully penetrant juvenile phenotypes (Fig. 2d and e). These phenotypes include alterations in neuropeptide expression patterns (e.g., NLP-45::T2A::GFP::H2B) that are directly regulated by LIN-14 and *lin-4* expression(*29*), as well as the perdurance of L1-stage expression programs (e.g., *oig-1* expression(*30*)) in later larval stages that antagonize the stage-specific rewiring of motor neuron synapses in the ventral nerve chord. Together, these results show that MYRF-1 binding sites upstream of *lin-4* are required across somatic lineages, and lineage-specific differences in miRNA transcriptional regulation reflect chromatin accessibility patterns rather than distinct timing mechanisms. Thus, organism-wide temporal coherence of heterochronic microRNA expression arises from global coordination of MYRF-1 acting on shared cis-regulatory elements, rather than from repeated convergence of tissue-restricted transcriptional programs.

### LIN-42 physically binds to MYRF-1

Gene regulatory networks that generate pulsatile transcriptional patterns typically rely on coupled negative feedback and delay mechanisms in which transcription factors activate their own repressors, producing periodic bursts of gene expression with defined phase, amplitude, and duration(*31*). Because MYRF-1 directly activates *lin-42* (Fig. 1A) and LIN-42 is predicted to dampen transcriptional pulses (*12, 13*), we asked whether LIN-42 physically associates with MYRF-1. Yeast two-hybrid (Y2H) assays showed that both major LIN-42 isoforms (LIN-42A and LIN-42B) robustly interact with the NDs of MYRF-1 and MYRF-2 (Fig. 3a), suggesting that these interactions may be part of a transcription/translational feedback loop (TTFL) mechanism. A hallmark feature of TTFL circuits is reciprocal buffering, in which an asymmetry in regulatory interactions caused by a loss-of-function mutation in one component can be suppressed by a loss-of-function mutation in the other (*32*). We therefore tested whether *myrf(lf*) and *lin-42(lf*) mutations modulate each other’s developmental timing phenotypes. The hypomorphic allele *myrf-1(mg412*) induces a supernumerary adult molting phenotype marked by aberrant *mlt-10p::GFP-pest* reactivation in adult animals (Fig. 3b)(*25*). Combining a *lin-42(lf*) allele with *myrf-1(mg412*) fully suppressed these defects (Fig. 3b and c). Conversely, the precocious heterochronic phenotypes associated with multiple *lin-42(lf*) alleles were eliminated when combined with *myrf-1(mg412*) (Fig. 3d and e; Table 1). *myrf-2(0*) mutations, which alone do not elicit detectable molting or temporal patterning phenotypes, also partially suppress precocious adult-alae formation observed in *lin-42(lf*) animals (Table 1). These reciprocal genetic interactions indicate that MYRF-1 and LIN-42 mutually regulate each other’s activity *in vivo* and support their placement in a shared feedback loop.

**Figure 3.**
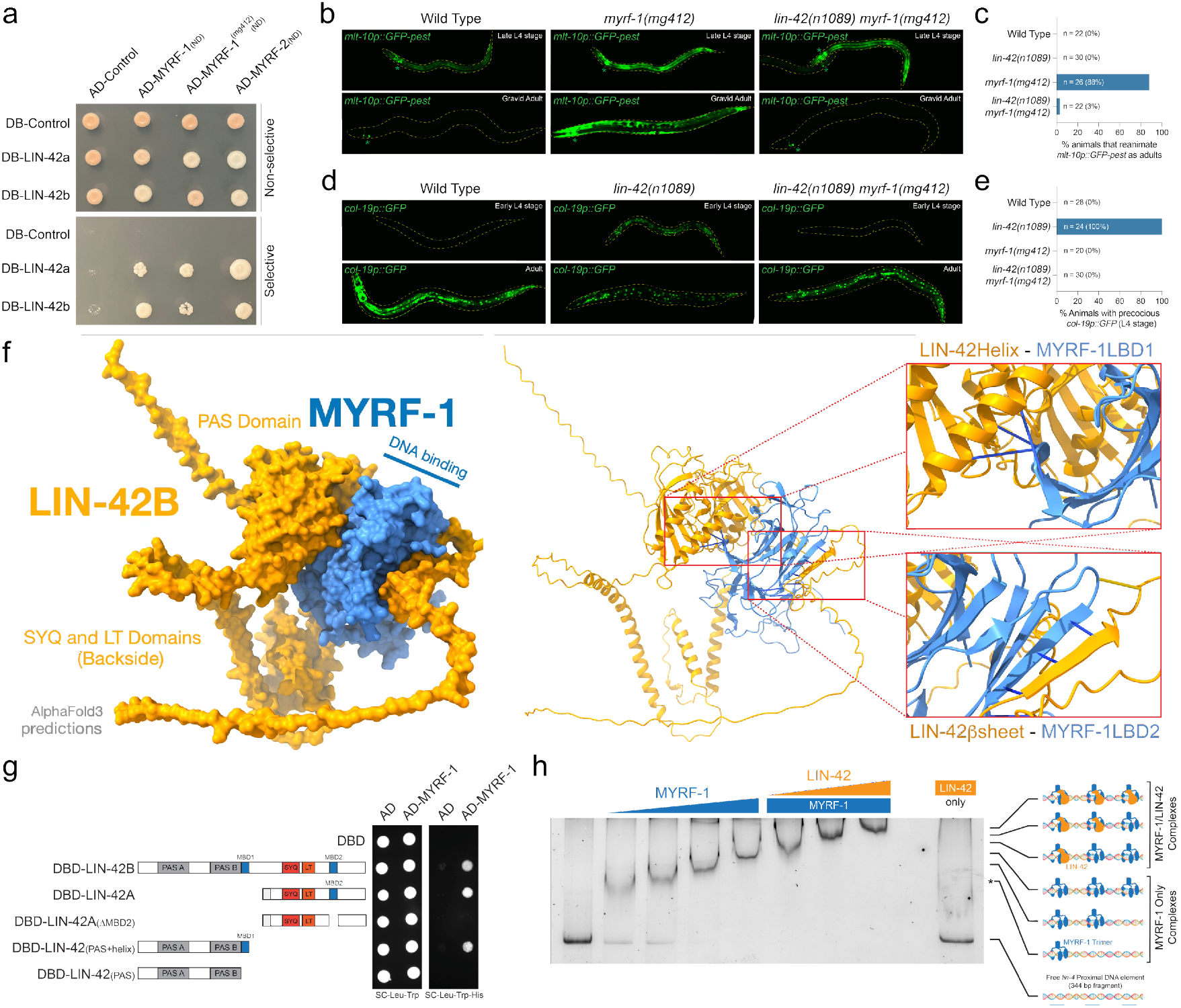
LIN-42 and MYRF-1 interact physically and genetically. **a**, Two-hybrid assays demonstrate that both isoforms of LIN-42 physically associate with the N-terminal nuclear fragment of the *C. elegans* MYRF proteins. Both LIN-42 isoforms also interact with MYRF-1(mg412). **b and c**, a hypomorphic allele of *myrf-1, myrf-1(mg412*), exhibits a highly penetrant supernumerary molting phenotype and reactivation of a *mlt-10p::GFP-pest* transcriptional reporter in adulthood. Combining a *lin-42(lf*) mutation with the *myrf-1(mg412*) allele suppresses these defects. **d and e**, Precocious expression of the *col-19p::GFP* transcriptional reporter in *lin-42(lf*) mutants is suppressed when combined with a *myrf-1(lf*) mutation. **f**, AlphaFold3 predictions of LIN-42 and MYRF-1 polypeptides predict that LIN-42 binds to two separate surfaces of a MYRF-1 monomer. The first AlphaFold3-predicted LIN-42 MYRF-1-binding Domain 1 (MBD1) comprises a conserved α-helical structure immediately C-terminal to the PAS domains. The second predicted interaction surface, MYRF-1-binding Domain 2 (MBD2), involves a β-strand of LIN-42 that extends an existing β-fold element in the predicted MYRF-1 structure. **g**, Two-hybrid experiments using LIN-42 constructs lacking amino acids implicated in MBD1 or MBD2 indicate that these regions of LIN-42 mediate the two-hybrid interactions. **h**, Increasing concentrations of a recombinant MYRF-1 result in a stepwise series of binding interactions with a proximal DNA region upstream of the *lin-4* gene, which harbors three predicted MYRF-1 consensus binding sites (Fig. 1a). This is the same DNA element, deleted in the *csh110* allele of *lin-4* outlined in Figure 2a, that harbors three predicted MYRF-1 trimer-binding sites. The addition of recombinant LIN-42B protein into the binding reactions results in three additional super shifts of the 3xMYRF-1/DNA complexes. LIN-42B alone exhibits no DNA-binding activity.

To define the molecular basis of this interaction, we used AlphaFold 3(*33*) to model complexes of MYRF-1(ND) and LIN-42. The predicted structures revealed two distinct LIN-42B domains that bind separate sites of MYRF-1(ND) (Fig. 3f). The first interface is mediated by a conserved α-helical extension immediately C-terminal to the LIN-42B PAS domain (MBD1), whereas the second involves a LIN-42B β-strand segment (MBD2) that intercalates into a β-fold domain of MYRF-1. To test these predictions, we generated LIN-42 deletion constructs lacking either the PAS-adjacent α-helix or the β-strand–forming segment and retested Y2H interactions between these proteins. Deletion of either segment from LIN-42 fragments abolished MYRF-1–LIN-42 association in yeast, whereas constructs retaining either motif preserved robust binding (Fig. 3g). These results support features of the structural predictions for this complex and indicate that specific LIN-42 domains are required for interaction with MYRF-1.

We next tested whether LIN-42 affects the ability of MYRF-1 to bind DNA by performing electrophoretic mobility shift assays with a long *lin-4* proximal DNA fragment deleted in the *csh110* allele of *lin-4*. This region harbors three predicted MYRF-1 consensus binding sites, which we speculate to each be bound by MYRF-1 homotrimers (Fig. 1a, 1e, and 2a). MYRF-1 titration revealed three discrete protein–DNA complexes consistent with sequential site occupancy of each predicted MYRF-1 binding site (Fig. 3h). We then titrated recombinant LIN-42B into these reactions and found that increasing LIN-42 concentrations led to a stepwise series of three additional super-shifts. Recombinant LIN-42B alone does not bind the *lin-4* proximal element (Fig. 3h), indicating that the super-shifts of the *lin-4* proximal element in these conditions result from LIN-42 binding to the three DNA-bound MYRF-1 complexes. This was supported by LIN-42B titration experiments with a single MYRF-1 binding site, which showed a single super-shifted species consistent with a 1:1 binding stoichiometry, as determined by mass photometry (Fig2 X).

### LIN-42 post-translationally controls the duration of MYRF-1 expression

To assess the functional importance of the LIN-42/MYRF-1 interaction domains *in vivo*, we used CRISPR/Cas9 editing to delete the two LIN-42 domains suspected to mediate direct MYRF-1 binding: the PAS-adjacent α-helical region (*lin-42(csh86)*) and the predicted, C-terminal β-strand segment (*lin-42(csh83)*). Animals lacking these individual domains showed normal larval cell division patterns, expressed *col-19p::GFP* only in adulthood, and developed adult alae at the correct time (Fig. 4a; Table 1). Strikingly, simultaneous deletion of both domains, *lin-42(csh83 csh86*), caused strong precocious phenotypes characteristic of significant *lin-42* loss of function, including early *col-19p::GFP* expression in L4-stage animals and the premature formation of adult-specific alae after the L3 molt (Fig. 4a; Table 1) (*13, 19*). These findings demonstrate that the two LIN-42 structural motifs function together *in vivo* and that most LIN-42 functions required for temporal cell-fate specification depend on its ability to interact with MYRF-1.

**Figure 4.**
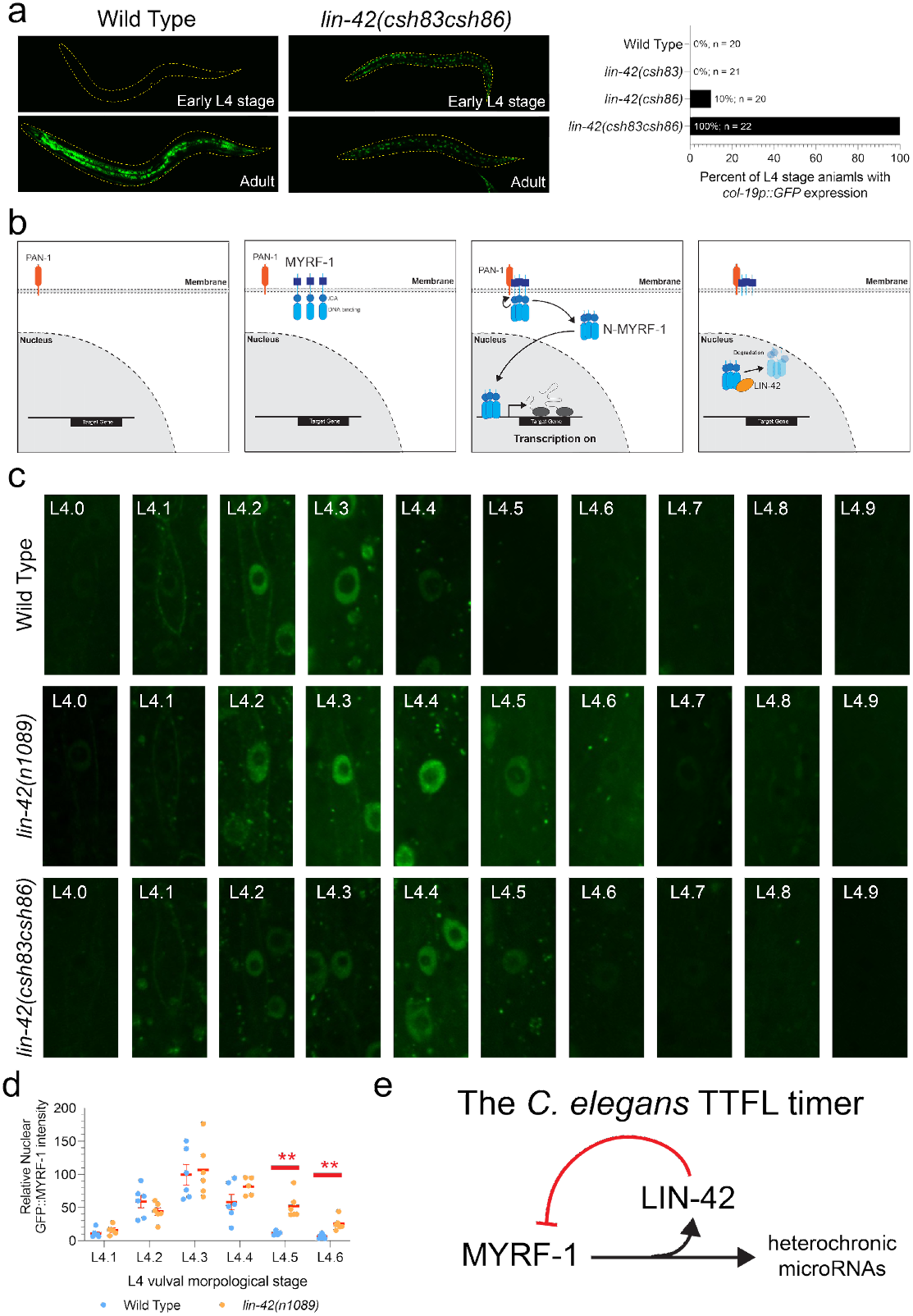
LIN-42 controls the duration of MYRF-1 nuclear residency to control temporal patterning. **a**, Simultaneous deletion of both MYRF-1 binding regions of LIN-42 results in strong heterochronic phenotypes, including the precocious expression of an adult-specific transcriptional reporter. **b**, MYRF-1 is initially translated in the cytoplasm and rapidly inserted into the ER and trafficked to the cytoplasmic membrane. Once concentrated on the membrane, the N-terminal fragment is autocatalytically cleaved and transported into the nucleus. In the nucleus, MYRF-1 binds to its regulatory elements upstream of its target genes to promote their transcription. **c**, Expression dynamics of GFP::MYRF-1 in hypodermal cells of L4-staged animals. **d**, Quantification of the dynamic changes in GFP::MYRF-1 expression in wild-type and *lin-42(n1089*) mutants. The perdurance of GFP::MYRF-1 in *lin-42(n1089*) mutants differs statistically from wild-type expression at the L4.5 and L4.6 stages using the Student’s t-test. ** indicates p = <0.01. **e**, A model depicting the regulatory interactions between MYRF-1, LIN-42, and miRNA target genes that compose a simple transcriptional/translational feedback loop.

Given the dynamic, once-per-larval-stage expression pattern of *myrf-1* (fig. S1a) and LIN-42’s role in repressing the transcription of MYRF-1 targets (Fig. 1a) (*12, 13, 21, 34*), we hypothesized that LIN-42 might post-translationally regulate MYRF-1 dynamics. To test this, we tracked GFP::MYRF-1 expression in L4-staged hypodermal cells, where GFP::MYRF-1 levels could be directly correlated with specific developmental milestones(*13, 35*). GFP::MYRF-1 was first observed on the outer membranes of lateral seam cells in early L4 animals (L4.0 stage) (Fig. 4c). Soon after initial detection in the membrane, cleaved GFP::MYRF-1(ND) started transitioning to the nucleus and was fully nuclear by the L4.3 stage (Fig. 4c). GFP::MYRF-1 levels then decreased starting at L4.4 and were absent from lateral seam cell nuclei by L4.6 stage. These dynamic expression patterns were also seen in other somatic cells (Fig. Sx). We examined GFP::MYRF-1 in animals with a large *lin-42* deletion allele, *lin-42(n1089*), or animals expressing the LIN-42 variant (LIN-42(csh83 csh86)) that cannot bind MYRF-1. In both *lin-42* mutant strains, early L4 GFP::MYRF-1 expression dynamics (membrane localization followed by nuclear import) remained unchanged (Fig. 4c). However, nuclear MYRF-1 expression dynamics are significantly altered in *lin-42* loss-of-function mutants, with nuclear MYRF-1 expression persisting in somatic cells for up to 1.5-2 extra hours (Fig. 4c; Fig. Sx)(*35*). The prolonged nuclear accumulation of MYRF-1 in *lin-42* mutants coincides with an extended period of *lin-4* transcriptional bursting in *lin-42* mutants(*13*), indicating that the timing of MYRF-1 nuclear activity determines the length—and likely the amplitude of *lin-4* transcriptional output.

## DISCUSSION

Our findings demonstrate that the reciprocal regulation between MYRF-1 and the PERIOD-like repressor LIN-42 forms a molecular timer that governs once-per-stage oscillations in gene expression. In this process, MYRF-1 directly activates *lin-42* transcription, while LIN-42 provides feedback to limit the duration of each MYRF-1 pulse. This transcriptional/translational feedback loop produces rhythmic MYRF-1 accumulation to precisely regulate the phase, amplitude, and duration of miRNA transcription. The observation that these pulses occur across somatic tissues and align with key developmental transitions suggests a mechanism in which the MYRF-1/LIN-42 timer coordinates gene expression timing with overall organism development. Furthermore, the fact that *myrf-1(0*) animals arrest immediately after molting indicates that this timer plays an essential role in promoting overall developmental progress, linking gene regulatory programs to the physiological processes necessary for transitioning between stages.

A critical feature of the developmental timer is that its activity within each larval stage is embedded in the nested-repression architecture of the heterochronic pathway(*9, 36, 37*). Each stage is defined by a temporal identity gene that both activates the stage-appropriate transcriptional program and represses the miRNAs expressed in the subsequent stage(*8, 38*). As the temporal identity gene declines late in the stage, repression of the corresponding miRNA loci is relieved, but transcription does not occur immediately. Derepression licenses transcriptional competence, whereas activation is imposed by the subsequent MYRF-1 pulse, which arrives only after the next cell fate has been specified. This delay ensures that miRNA expression follows, rather than precedes, fate commitment, and the resulting miRNA expression wave in the subsequent stage represses the next temporal identity gene, converting rhythmic MYRF-1/LIN-42 activity into irreversible developmental transitions.

These results identify a developmental timing circuit that synchronizes gene expression across tissues by coupling rhythmic transcriptional activity to irreversible fate transitions. Although the MYRF-1/LIN-42 timer shares transcriptional/translational feedback logic with circadian clocks(*31, 39*), it serves a distinct function by scheduling a finite series of sequential events that occur once and must be executed in the correct order. By interfacing with the nested-repression architecture of the heterochronic pathway, this timer converts oscillatory gene expression into stage-locked miRNA waves that drive unidirectional developmental progression. Together, these findings show how conserved clock components can be repurposed to coordinate organism-wide development and couple gene regulatory dynamics to growth.

## Acknowledgments

ChatGPT v5.1 was used for polishing initial drafts.

## Author contributions

Methodology: PW, JW, BP, CMH

Investigation: PW, JW, BP, IV, DR, KL, JT, CMH

Visualization: CMH

Funding acquisition: CMH, LJ, JK

Project administration: CMH Supervision: CMH, LJ

Writing – original draft: CMH

Writing – review & editing: PW, BP, IV, JW, LJ, CMH

## Data and materials availability

All data, code, and materials used in the analysis are available to any researcher for purposes of reproducing or extending the analysis.

## MATERIALS AND METHODS

### *C. elegans* strains and maintenance

*C. elegans* strains were maintained on standard nematode growth medium (NGM) plates seeded with *E. coli* OP50 at 20 °C or 15°C under standard laboratory conditions. The Bristol N2 isolate was used as the wild-type reference strain. A complete list of strains used in this study is provided in Supplemental Table SX.

### CRISPR genome deletion and GFP tagging

Genome editing and endogenous GFP tagging were performed by following established CRISPR/Cas9 protocols(*40, 41*). Briefly, Cas9–sgRNA ribonucleoprotein (RNP) complexes were preassembled by combining purified recombinant Cas9 protein with synthetic CRISPR sgRNAs targeting specific genomic loci, together with a *dpy-10* sgRNA used as a co-CRISPR marker to generate Roller phenotypes. The injection mixture, containing the assembled RNP complexes and a PCR-amplified repair template with flanking homology arms, was injected into the germline of hermaphrodites. Broods segregating Roller progeny were screened for genome edits, and putative transgenic animals were subsequently genotyped by PCR to confirm domain deletions within the *lin-42* gene, MYRF-1 binding sequences upstream of *lin-4*, or endogenous GFP tagging at the *myrf-1* N terminus.

### Yeast two-hybrid assays

Plasmids encoding target proteins fused to GAL4 DNA-binding-domain (pBD) and GAL4 Activation Domain (pAD) were co-transformed into the pJ69-4a Y2H yeast strain (*42*) using the lithium acetate method as previously described in the Matchmaker™ GAL4 Two-Hybrid System 3 User Manual (Takara Bio USA, Inc.). Transformants were selected on SC-TRP-LEU plates for 3 days at 30 °C. Three independent colonies from each transformation were subsequently spotted onto SC–HIS–TRP–LEU plates. Protein-protein interactions were inferred from visible growth on 3-AT conditions with negative growth in empty vector controls after 3 days of incubation at 30 °C.

### ChIP-seq

Endogenously GFP-tagged MYRF-1 animals were synchronized at the L1 stage by hypochlorite treatment of gravid adults followed by overnight hatching in M9 buffer. Synchronized L1 larvae were plated on 150-mm NGM plates seeded with *E. coli* OP50 and grown for ∼11 h, when GFP::MYRF-1 is predominantly localized to the nucleus. Approximately 100 µL of packed worms were collected by washing in M9 buffer, crosslinked in 2% formaldehyde, and quenched with 125 mM glycine. Two biological replicates were collected for each ChIP experiment. ChIP–seq was performed as previously described (*13*). Briefly, crosslinked animals were homogenized in FA buffer (50 mM HEPES–KOH pH 7.5, 150 mM NaCl, 1 mM EDTA, 1% Triton X-100, 0.1% sodium deoxycholate, and 0.1% sarkosyl) supplemented with protease inhibitors, dounce-homogenized on ice, and sonicated at 4 °C to shear DNA and generate 200–800 bp chromatin fragments. Clarified extracts were quantified by Bradford assays, and 1–4 mg of total protein was incubated overnight at 4 °C with anti-GFP antibody (Abcam, #ab290) and anti-RNA polymerase II antibody(Millipore 05-952-I-100UG). Immune complexes were captured with protein A/G Sepharose beads, sequentially washed with buffers of increasing stringency, and eluted in SDS-containing ChIP elution buffer. Crosslinks were reversed overnight at 65 °C, followed by RNase A and Proteinase K treatments. DNA was purified using Qiagen MinElute columns and analyzed for fragment size before library preparation and sequencing with Illumina NextSeq500 at NYU Center for Genomics and Systems Biology core facility.

### ChIP-seq data analysis and MYRF-1 binding motif identification

Single-end 75bp ChIP and input control reads were processed using established computational pipelines. Raw sequences were first assessed for quality with FastQC and FastQ Screen, and low-quality reads were removed using Trimmomatic. Adapter sequences were trimmed with Cutadapt, and the filtered reads were aligned to the *C. elegans* reference genome (ce11) using Bowtie2(*43*). MACS2 (*44, 45*) was used for peak calling, with input DNA as the control, and significance thresholds of P < 0.001 and q < 0.05 were applied. Reproducible peaks across biological replicates were identified by intersecting using BedTools (*46*). A random set of genomic regions matched in number and length to the reproducible peaks was generated as background controls. Peak and control sequences were analyzed with the MEME suite in discriminative mode using a first-order (dinucleotide) background model. Two enriched motifs were identified: a predominant GA-repeat motif (dimeric or trimeric) present in most peaks, and a less repetitive secondary motif found in a smaller subset. These motifs were scanned across the peak and control sequences using FIMO(*47*), and the resulting P-values of the top hits were used to generate ROC curves. The GA-repeat motif achieved an area under the curve (AUC) of 0.78, and the secondary motif, 0.73. Based on these curves, motif significance thresholds were set at P < 1 × 10^−4^ and P < 1 × 10^−3^ for the GA-repeat and secondary motifs, respectively. Genome-wide motif scanning was then performed using these thresholds, and the resulting motif distributions were visualized as custom UCSC Genome Browser tracks.

### Confocal imaging

For confocal imaging, worms at the appropriate developmental stages were mounted on 2% (w/v) agarose pads in 100 mM levamisole (Sigma). Images were acquired using a Hamamatsu Orca EM-CCD camera and a Borealis-modified Yokagawa CSU-10 spinning disk confocal microscope (Nobska Imaging, Inc.) with a Plan-APOCHROMAT x 100/1.4, 63x/1.3, or 40/1.4 oil DIC objective controlled by Visiview Software (version: 7.0). LED illumination at 488 nm and 561 nm was used to excite green and red fluorophores, respectively. Images were processed in ImageJ (Fiji) using identical processing settings for all genotypes and developmental stages within each experiment.

### Quantification of fluorescent reporter

The average intensity (arbitrary units) of GFP::MYRF-1 in hypodermal cells of L4-stage animals was quantified using ImageJ as previously described (*13, 17*). For each cell, fluorescence intensity was calculated as the nuclear signal minus the background signal measured from the same image. The mean intensity of three hypodermal cells was used to determine the GFP::MYRF-1 level for each animal. All statistical analyses were performed using GraphPad Prism 9 (GraphPad Software, San Diego, Ca). Mean ± SEM values were calculated and plotted in Prism. Differences between the two groups were considered statistically significant when p < 0.05 (Student’s t-test).

### Recombinant protein expression and purification

*C. elegans* MYRF-1(ND)(amino acids 1-483) and full-length LIN-42B were cloned as N-terminal Strep-SUMO fusion proteins in separate pFL vectors of the MultiBac Baculovirus expression system(*48*). These proteins were individually expressed in Sf9 cells grown in CCM3 media (Hy-Clone) at 27°C for 60 hr. Cells were pelleted by centrifugation at 1800 rpm for 20 min, resuspended in lysis buffer (MYRF-1(ND) = 25 mM Tris pH 8, 200 mM NaCl, 2 mM DTT, 10% glycerol; LIN-42B = 20 mM Tris pH 8, 200 mM NaCl, 5 mM BME) and sonicated in the presence of homemade protease inhibitor cocktail and cOmplete mini EDTA-free protease inhibitors (Roche). Lysates were clarified by ultracentrifugation for 1 hr at 4°C. For affinity chromatography, lysate supernatants were batch-bound to Strep-Tactin Superflow resin (IBA) for 1 hr at 4°C with rotation. Affinity beads were harvested by centrifugation at 1000 rpm for 3 min, decanted to 1 column volume (CV), resuspended, and applied to a gravity column. For MYRF-1(ND), the column was washed with 1 CV lysis buffer, 2 CV high salt wash buffer (25 mM Tris pH 8, 500 mM NaCl, 2 mM DTT, 10% glycerol), and 1 CV lysis buffer to remove nucleic acid and protein contaminants. For LIN-42B, the column was washed with 3 CV lysis buffer. Proteins were eluted with 5 mM desthiobiotin in lysis buffer. The strep-SUMO tag was removed from MYRF-1(ND) by TEV protease (1:30 by mass) overnight at 4ºC. Proteins were further purified using anion exchange (HiTrap Q at pH 8) and size exclusion chromatography (Superose 6 increase, 10/300) in storage buffer (25 mM Tris pH 8, 200 mM NaCl, 2 mM DTT, 10% glycerol). Peak fractions were assessed for purity by SDS-PAGE, pooled, and concentrated to 0.6 mg/mL (MYRF-1(ND)) or 0.9 mg/mL (LIN-42B). To reconstitute the MYRF-1(ND) trimer onto DNA, fully purified MYRF-1(ND) was incubated with annealed proxB DNA at a 5-molar excess in MYRF-1(ND) storage buffer on ice for 1 hr. The DNA-bound trimer was separated by analytical gel filtration on a Superose 6 increase 3.2/300 in MYRF1(ND) storage buffer. Peak fractions were analyzed by SDS-PAGE, and gels were stained with SYBR gold (1:33k in ddH2O) at room temperature for 10 min to highlight co-purified DNA.

### Electrophoretic mobility shift assays

For competitive EMSAs, MYRF-1(ND) was incubated with ATTO 680-labeled proxB on ice for 5 min, after which competitor oligos were added. Reactions were incubated with competitors for an additional 15 min at room temperature and run on 5% TBE gels in 0.5x TBE at 135V for 35 min at 4ºC. For LIN-42B titration experiments, MYRF-1(ND) was incubated with unlabeled proxB (44 bp) or a *lin-4* proximal promoter fragment (344 bp) in the presence of serially diluted strep-SUMO-LIN-42B for 15 min at room temperature. Reactions with proxB were run on 5% TBE gels in 0.5x TBE at 110V for 45 min, while those with promoter *lin-4* proximal were run at 100V for 90 min. All reactions were conducted under conditions similar to those used in Ndt80 EMSAs (10 mM Tris pH 8, 75 mM KCl, 11 mM MgCl2, 50 uM ZnSO4, 10% glycerol, 1 mM DTT, and 0.02% Tween-20), and all gels were prerun at 100V for 30 min at 4ºC. Experiments with unlabeled probes were stained with a 1:100k dilution of SYBR gold in ice-cold 0.5x TBE for 3 min and destained for 5 min prior to imaging.

### Mass photometry

MYRF-1(ND) was diluted to 1.2 uM in storage buffer and incubated with or without prox B at a 5:1 molar ratio in reaction buffer (25 mM Tris pH 8, 150 mM NaCl, 2 mM DTT, 10% glycerol). Reactions with DNA were incubated for 15 min at room temperature prior to taking measurements, while those without were measured immediately after dilution. Reconstitution of the MYRF-1(ND):LIN-42B:DNA ternary complex was performed in a similar way, this time incubating proxB DNA with a 5-molar excess of each of MYRF-1(ND) and strep-SUMO-LIN-42B for 15 min at room temperature. Strep-SUMO-LIN-42B control samples were prepared at 6 nM in storage buffer and immediately analyzed after dilution. Samples were analyzed at 1/10 the indicated prepared concentration in 1xPBS on untreated MassGlass UC slides (Reyfeyn). Movies were recorded for 60 sec in AcquireMP, and data were analyzed in DiscoverMP software (Reyfeyn). Beta-amylase derived from sweet potato was used to create the standard curve.

